# Hepatoblastomas with carcinoma features represent a biological spectrum of aggressive neoplasms in children and young adults

**DOI:** 10.1101/2021.07.15.445600

**Authors:** Pavel Sumazin, Tricia L. Peters, Stephen F. Sarabia, Hyunjae R. Kim, Martin Urbicain, Emporia Faith Hollingsworth, Karla R. Alvarez, Cintia R. Perez, Alice Pozza, Mohammad Javad Najaf Panah, Jessica L. Elswood, Kathy Scorsone, Howard Katzenstein, Allison O’Neal, Rebecka Meyers, Greg Tiao, Jim Geller, Sarangarajan Ranganathan, Arun A. Rangaswami, Sarah E. Woodfield, John A. Goss, Sanjeev A. Vasudevan, Andras Heczey, Angshumoy Roy, Kevin E. Fisher, Rita Alaggio, Kalyani R. Patel, Milton J. Finegold, Dolores H. López-Terrada

## Abstract

Malignant hepatocellular cancers are the most common primary liver malignancies in children, and hepatoblastomas (HBs) account for more than two-thirds of these cases. While most HBs respond to chemotherapy and have favorable outcomes, the 3-year overall survival rate for high-risk HBs is below 50% and guidelines for their classification and treatment are still evolving. HB risk-stratification efforts using clinical, histological, and molecular parameters have been reported to help identify patients that require more or less aggressive therapies in retrospective studies, and are being validated in clinical trials. However, risk assessment is particularly challenging for cancers with certain histologies, including tumors in the recently proposed provisional *hepatocellular neoplasm not otherwise specified* (HCN NOS) category. HCN NOSs exhibit either intermediate or combined HB and hepatocellular carcinoma (HCC) histological features, and while neoplasms with such features were observed over a decade ago, only a handful have been characterized and little is known about their biology and clinical features. Here, we molecularly characterized a series of clinically annotated HCN NOSs that demonstrated either intermediate HB/HCC histology or distinct coexisting areas with HB and HCC histological features. In addition, molecular profiling of HBs demonstrating focal pleomorphism or anaplasia (HB FPA) revealed underlying biological features previously observed in HCCs. Our study suggested that HCN NOSs and HB FPAs are aggressive tumors, irrespective of patient age or resectability. Consequently, we designated them collectively as *HBs with carcinoma features* (HBCs) and outlined histological and molecular characteristics for their diagnosis and treatment. In our single-institution study, transplanted HBC patients were significantly and more than twice as likely to have good outcomes, highlighting the importance of molecular testing and aggressive early intervention.

## INTRODUCTION

Hepatoblastomas (HBs) and hepatocellular carcinomas (HCCs) are the most common primary liver malignancies in children. HBs are usually diagnosed in young children and have shown the highest average annual percent increase in incidence over past decades (1). HCCs are more commonly seen in older children and adolescents and are often associated with underlying genetic, metabolic, and inflammatory conditions (1). HB- and HCC-clinical presentations, treatment options, and outcomes differ dramatically, with 5-year overall survival rates near 70% for HB and 30% for HCC patients (2).

HBs are embryonal neoplasms that histologically recapitulate stages of liver development (3). They may be either exclusively epithelial or mixed epithelial and mesenchymal, and are characterized by aberrant WNT-pathway activation that is most often associated with genetic alterations of *CTNNB1* (4). These cancers often respond to chemotherapy, and a combination of cisplatin and doxorubicin-based chemotherapy regimens and surgery are an effective treatment for most patients. However, incomplete response to chemotherapy often leads to unfavorable clinical outcomes (5). Pediatric HCCs are a histologically and biologically diverse group of neoplasms (6), when compared to HBs, and are generally clinically aggressive, chemotherapy-resistant tumors, with complete surgical resection being their only cure (2).

Recently, international efforts to characterize the histopathology of pediatric hepatocellular malignancies and create a consensus classification to be used in collaborative international trials, recognized the existence of a difficult-to-diagnose histological group of hepatocellular tumors in children and young adults demonstrating either combined or overlapping histological features of HB and HCC (7). The provisional HCN NOS diagnostic category was proposed to identify and further characterize these difficult to classify tumors. Although tumors with HCN NOS features were first documented over a decade ago (8, 9), only recently a total of 5 specimens have been molecularly characterized (4, 10, 11). Results from these studies suggested that pediatric hepatocellular neoplasms may represent a broad histological and biological spectrum, that HCN NOS may carry characteristic alterations—including *TERT* promoter mutations—and that their prognosis may be particularly poor (4). We present a study of the outcomes and molecular features of HCN NOSs in addition to other HBs, including HBs with focal pleomorphism or anaplasia (HB FPAs). While the molecular and histological features of HB PFAs are present at diagnosis, today, focal pleomorphisms are often observed following treatment by chemotherapy and these tumors are often identified as HBs with pleomorphism in surgical pathology reports. We emphasize that the prognostic significance of these histological features and the underlying biology of HB PFAs remain unknown and do not inform current risk stratification protocols.

Current protocols for diagnosing and classifying pediatric hepatocellular neoplasms and their subtypes are based on histopathology and limited immunohistochemistry of diagnostic biopsy or resection specimens, which sometimes may be insufficient for definitive differentiation between HBs, HCN NOSs, and HCCs (8). In particular, HCN NOS tumors have high intra-tumor heterogeneity (8, 10) and their classification, when based on needle biopsies, can be particularly challenging. Here, we analyze the molecular and genetic features, as well as treatment and outcomes data, of the largest cohort of histologically confirmed HCN NOSs. We propose that risk stratification based on a combination of histology and molecular features, which have been key to increasing cure rates for other pediatric cancers, will help in recognizing, characterizing, and treating these and other pediatric hepatocellular neoplasms. Based on common molecular features and outcomes we propose the new designation hepatoblastoma with carcinoma features (HBC), which includes pediatric hepatocellular neoplasms in the HCN NOSs, and HB FPAs—all with molecular features reported in HCCs. We note that while histological and molecular features HB FPAs can be identified at diagnosis, today, focal pleomorphism and anaplasia are more often seen in post-chemotherapy resection specimens. Finally, our retrospective analysis of single-institution data suggested that while stages 2 and 3 HBCs, as scored according to the Pretreatment Extent of disease staging system (PRETEXT), had poor outcomes when treated by chemotherapy and resection alone, transplantation significantly improved their outcomes.

## EXPERIMENTAL PROCEDURES

### Case selection

With institutional review board approval, we searched the Texas Children’s Hospital pathology database to identify in-house HB and HCC cases, as well as cases that were seen in consultation between 2004 and 2016. From 170 cases, we identified a total of 25 HCN NOS cases, and 10 HB FPAs, that were diagnosed in patients that were younger than 8 years at diagnosis. The age criterion for HB PFAs was intended to distinguish the biology and outcomes of patients with these tumors from those with HBs without these features that were diagnosed in children older than 8 years—these older patients are assigned a higher risk in current clinical trials (12, 13) and we aimed to investigate the outcomes of HB PFAs that are not necessarily classified as high risk. In addition, 5 HB cases diagnosed in older children (greater than 8 years) and with unambiguous classic HB histology with typical epithelial or epithelial and mesenchymal HB features and without any atypical histological features, were selected as high-risk-HB controls. These samples were selected to test whether their molecular and genetic features better resemble those of HBs observed in younger children, HBCs, or HCCs. We note that 4 of the 5 older HB cases were transplanted and all but 1 had metastatic disease (Figure 2A). Cases were selected based on age, diagnoses in surgical pathology reports, histological reviews, and materials available for genetic testing. HCN NOS cases were identified based on entries in their diagnostic report which included a final diagnosis, or a diagnostic comment of HCN NOS or Transitional Cell Tumor. Similarly, HB FPAs were chosen to include patients whose diagnostic reports mentioned atypical histological features such as focal anaplasia, macrotrabecular pattern, or significantly increased pleomorphism. Pleomorphic components were defined as large (>2-3x) tumor cells with large hyperchromatic nuclei, prominent nucleoli, irregular nuclear contours, intranuclear inclusions, with or without atypical mitoses. Histological reviews of representative glass slides from all cases were performed by three pathologists (MJF, DLT, and TLP) who confirmed the features in the original reports and selected representative areas of tumor most suitable for molecular testing. Selected clinical information was collected for these patients, as allowed by the IRB protocol; see Table S1.

### Genetic profiling

Tumor genome-wide copy number was profiled using genomic DNA isolated from formalin-fixed paraffin-embedded (FFPE) tumor tissue (38 cases) using the Affymetrix OncoScan FFPE Assay Kit, according to the manufacturer’s protocol. CEL files generated from the array were analyzed using the OncoScan Console 1.3 software to create OSCHP files containing copy number calls. Data analysis was performed using the Affymetrix CHAS 3.1 software and OncoScan Nexus Copy Number 7.5 software and aligned to the National Center for Biotechnology Information (NCBI) human build GRCh37/hg19 assembly. In addition, 37 cases were interrogated using a next-generation sequencing assay designed to detect the presence or absence of mutations in coding and splicing regions of 2247 exons in 124 genes as well as the promoter region of *TERT*. Barcoded DNA libraries were constructed from genomic DNA samples using KAPA HyperPlus Kit. Capture hybridization-based target enrichment was performed using a custom-designed Roche Nimblegen SeqCap probe set, and was followed by sequencing on the Illumina® MiSeq® System. The FASTQ sequencing files generated were used for read alignment using the NextGENe v2.4.1.2 program, as well as the Burrows-Wheeler Aligner (BWA). Variant calling and annotation were performed by NextGENe and Platypus variant callers to produce a merged output of annotated variants. A literature review was then performed to determine the significance of each variant identified. To verify capture assay results, *CTNNB1* exon 3/4, *NFE2L2* exon 2, and *TERT* promoter mutation analysis were performed as previously described (4). For cases without viable RNA for *CTNNB1* RT-PCR mutation analysis, the same primer set was utilized with genomic DNA for long PCR mutation analysis (4). Predicted somatic mutations and CNAs are listed in Tables S2 and S3, respectively.

### RNA expression profiling

Gene expression profiling FFPE tumor tissues from 37 HBs was performed using the NanoString nCounter PanCancer Pathways Panel, which includes probes against 730 cancer genes, plus 30 HB-specific genes. The 30 custom selected probes added to the panel were selected based on differential expression observed in previously reported HB expression profiling efforts and to represent key hepatocarcinogenesis pathways (see panel design and normalized expression values in Table S4). Multiplexed measurements of gene expression through digital readouts of the relative abundance of mRNA transcripts were performed using the following steps: hybridization of RNA to fluorescent Reporter Probes and Capture Probes, purification of the target/probe complexes using nCounter Prep Plates and nCounter Cartridges on the nCounter Prep Station, and analysis using the nCounter Digital Analyzer to provide a test result. Gene expression profiles across samples were normalized to equate positive control probes and expression estimates were z-score normalized for each probe independently. To perform profile RNAs in single HB cells, we dissociated tumor cells from a fresh primary HB tumor using an enzyme mix in a gentleMACS C Tube (Miltenyi Biotec). Cells were spun, strained, and washed in PBS (Life Technologies) and BSA 0.04% (VWR), pooled, and loaded into the Chromium Next GEM Single Cell 5’ Library and Gel Bead Kit version 1.1 workflow (10× Genomics). Single-cell RNA and cell hashing library preparation was performed according to the manufacturer’s instructions. Libraries were sequenced by novoGene using the NovaSeq 6000 System and an S4 2 ×150 kit (Illumina).

## RESULTS

To investigate the histological, molecular, and clinical features of HCN NOSs and HB FPAs tumors (HBCs), we compared outcomes for HBCs and HBs diagnosed in patients older than 8, which have been reported to be high-risk (14). Our analyses of HBCs highlighted the dysregulation of and recurrent genetic alterations in cancer genes and pathways and suggested biological and clinical similarities between HBC subtypes by contrasting HBC molecular and clinical features with those that characterize epithelial and epithelial/mesenchymal HBs, and HCCs.

### Tumor histopathology and patient selection

Histological review of over 170 candidate cases with diagnostic reports at Texas Children’s Hospital identified a total of 25 HCN NOS and 10 HB FPAs in patients younger than 8 years old at resection. HB FPAs were selected with the age cutoff to distinguish them from those diagnosed in older patients (>8 years), which are considered to be associated with high risk according to The Children’s Hepatic tumors International Collaboration stratification algorithm CHIC-HS (14). We studied the biology and outcomes of HBCs and compared their histological features with those of lower-risk HBs, higher-risk HBs diagnosed in patients older than 8-years of age, and HCCs.

HCN NOSs were characterized by concurrent HB and HCC histological features and were classified into *biphasic HCN NOSs* and *equivocal HCN NOSs* (Figure 1). Biphasic HCN NOSs had discernable tumor areas with either distinct HB or HCC features, while equivocal HCN NOSs were characterized by cancer cells and growth patterns that were intermediate between HB and HCC, and, hence, difficult to classify (n=13). Biphasic HCN NOSs were distinguished from equivocal HCN NOSs in their presentation of homogenous regions and histological variability across regions, with some regions populated by cells with HB features and others with HCC features. In comparison, as shown in Figure 1, equivocal HCN NOSs were histologically homogeneous tumors overall with intermediate HB and HCC features, including higher NC ratios, occasional pleomorphism, increased mitotic count, and occasional macrotrabecular patterns (n=12). Some of these tumors had been previously classified as *transitional liver cell tumors*, a designation that has been replaced in the international consensus classification by HCN NOS^7^.

**Figure 1.**
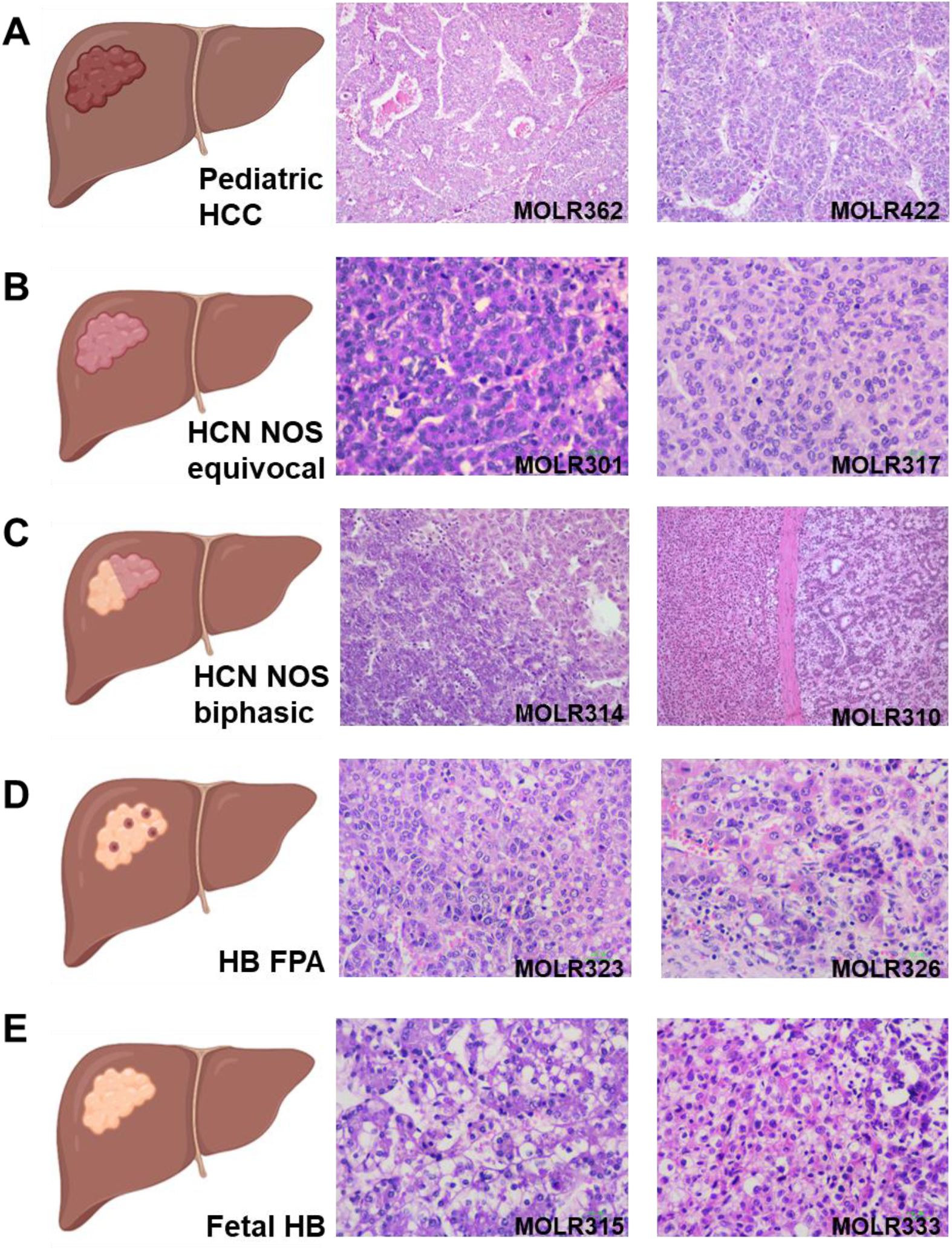
Representative histopathology of the pediatric hepatocellular neoplasms in the study. (**A**) Representative pediatric HCCs (4X). (**B**) Two equivocal HCN NOSs exhibiting homogeneous cell patterns with higher nuclear-to-cytoplasmic ratios, slight pleomorphism, increased mitotic count (10X). Equivocal HCN NOSs may display a macro-trabecular growth pattern (not illustrated). (**C**) Biphasic HCN NOSs exhibiting distinct histological patterns in separate nodules. Areas of typical HBs adjacent to HBs with carcinomatous features can be distinctly identified (4X). (**D**) HB FPAs showing histological features of HBs but containing foci or cell clusters with atypical cytologic features including increased pleomorphism, vesicular chromatin with prominent nucleoli, and higher nuclear-to-cytoplasmic ratio (10X). (**E**) Epithelial fetal HBs without atypical or carcinomatous features (10X).

In addition to HCN NOSs, we analyzed the biology and outcomes of 10 HB FPAs. These tumors focally displayed cells that featured increased pleomorphism, prominent nucleoli, and higher NC ratios, occasionally arranged in a macro-trabecular pattern (Figure S1). These are features that are most often seen in post-chemotherapy specimens and their underlying biology and prognostic significance remain unknown. However, HB FPAs can be recognized in diagnostic specimens pre-chemotherapy based on the combination of histological features and molecular features described in the following sections.

### HBCs are high-risk liver cancers

We defined good patient outcomes as event-free survival for at least 2 years, and poor patient outcomes as relapse or death of disease during a median follow-up of 5.5 years. Of the 25 HCN NOS patients, treatment data were available for 23; of which 16 underwent chemotherapy followed by resection and 7 underwent transplantation. Of the 10 HB FPA patients, treatment data were available for 9, of which 7 had chemotherapy followed by resection and 2 underwent transplantation. Of the 5 older patients with classic HB, 1 had chemotherapy followed by resection and 4 underwent transplantation (Fig 2B). Of the 40 patients, PRETEXT data were available for 34, of which 12 were Pretext 4, 18 Pretext 3, 2 Pretest 2, and 2 Pretext 1. Most (n=30, 88%) of our selected patients had PRETEXT 3 and 4 disease at diagnosis, where no two adjoining liver sections were free of disease; see Figure 2A. While feasible, complete resections of these cancers are more challenging and may depend on their response to preoperative chemotherapy (15, 16). While the majority (60%) of our HBC patients had poor outcomes, all patients with PRETEXT stages 1 and 2 cancers (n=4), which are expected to be fully resected, had good outcomes. Consequently, outcome-predictive features were most applicable for classifying PRETEXT stages 3 and 4 patients. Not surprisingly, liver transplantation (17, 18) was significantly predictive of improved patient outcomes in our single-institution cohort with a hazard ratio of 3.2 and a 95% confidence interval of 2.5 to 4.3, assuming a normal sampling distribution. In total, 7/9 (78%) of our transplanted HBC patients had good outcomes. By comparison, the presence of metastases, which is known to correlate with patient outcomes (19), had a hazard ratio of 1.8 and a 95% confidence interval of 1.3 to 2.4 and was not significantly predictive of outcomes in our patient cohort, likely to due to its small size (Figure 2B). Moreover, metastases detection did not reduce the benefit of transplantations, and transplanted patients had improved outcomes (p<1E-4) after conditioning on metastatic disease. We note that Figure 2B tables, which examined the associations between outcomes and metastatic disease or transplantation, included both HBCs older HB patients, resulting in significance estimates of p=0.07 and p=0.01, respectively. Excluding older patients resulted in significance estimates of p=0.06 and p=0.01, respectively. In addition, because of the risk and age biases in our selection criteria, age at diagnosis was not predictive of patient outcomes.

**Figure 2.**
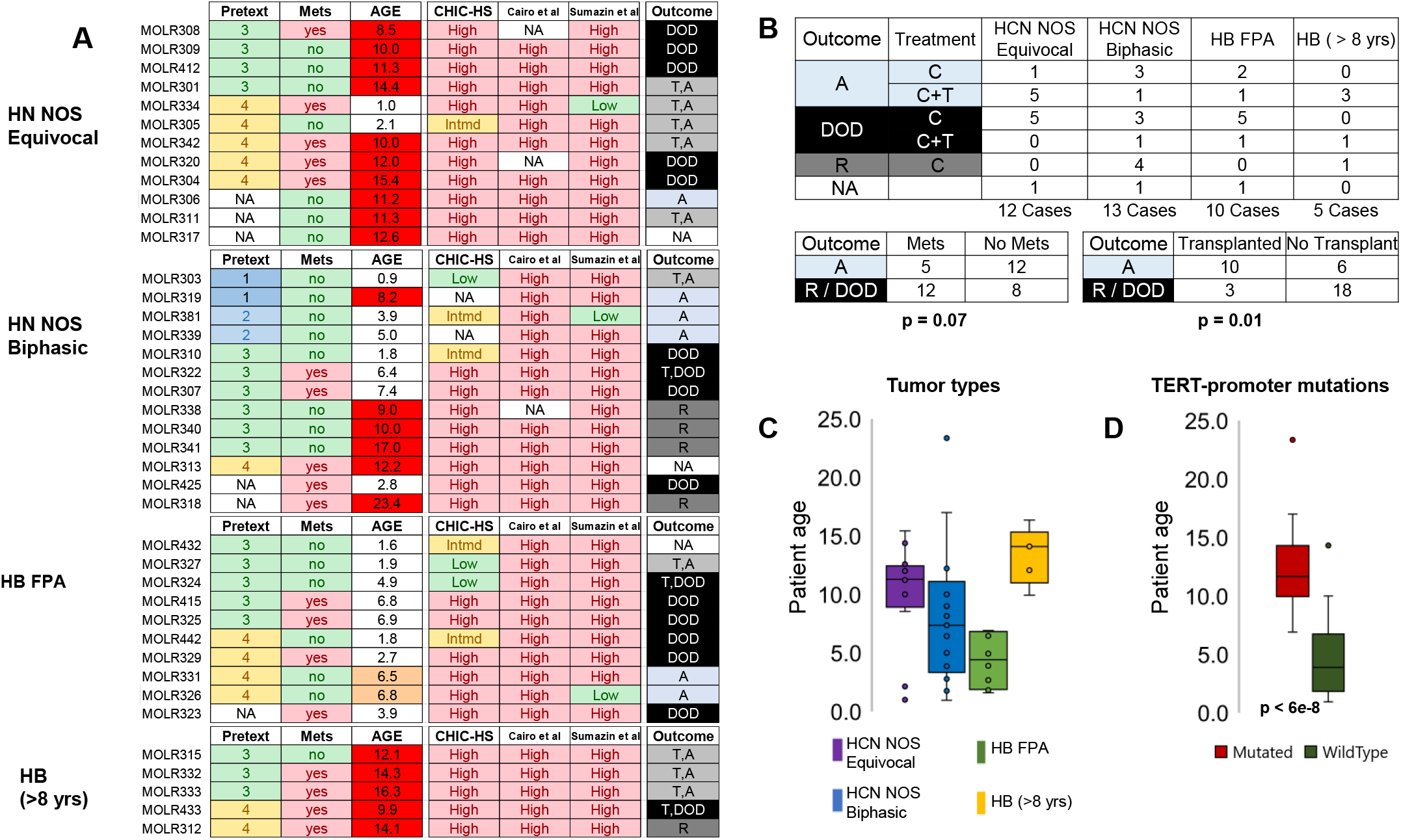
We evaluated the pathology of 40 HBs, including (**A**) 12 equivocal HCN NOSs, 13 biphasic HCN NOSs, 10 HB FPAs that were younger than 8-years old, and 5 classical HBs of patients of age greater than 8. Patient outcomes include alive with no evidence for recurrence for at least 2 years (A), dead of disease (DOD), relapsed (R), or not available (NA). Treatment strategies included chemotherapy (C) and transplantation (T). Available data for some of these patients include pretext scores, the presence of metastatic lesions, age at diagnosis, risk classification by CHIC-HS, Cairo et al., and Sumazin et al., as well as outcomes; note that the high-risk HB class C2 by Cairo et al. was marked as high-risk for readability. (**B**) HBC patients had worse outcomes than our older patient cohort (hazard ratio of 1.5). The presence of metastases correlated with outcomes and had a hazard ratio of 1.8, p<0.07. Transplanted patients had significantly better outcomes with a hazard ratio >3 and p<0.01. Both transplanted patients with and without detected metastases had improved outcomes with a hazard ratio >3 and p<1E-4 after conditioning on metastatic disease. Note that patients without available outcomes data were excluded from these comparisons. (**C**) Our HB patients differed by age, with most equivocal NOS patients older than most biphasic NOS patients. (**D**) *TERT*-promoter mutations were mainly observed in tumors diagnosed in older patients with no significant associatiations with other tumor classifications.

Outcomes for older HB patients are generally expected to be poor (14), and patients with PRETEXT stages 3 and 4 HBs that are older than 8 years of age at the time of diagnosis are uniformly considered high risk and are treated aggressively, including by early liver transplantation for unresectable tumors. We note that the age composition of our cohort was not balanced. Equivocal HCN NOS patients were older, on average, than biphasic HCN NOS patients, and the other two subtypes—HB FPAs and older HBs—were younger and older, respectively; see Figure 2C. Our set of older HBs (older than 8 years at diagnosis) with classical histologies included 5 patients: 3 (60%) good- and 2 (40%) poor-outcome patients. All 3 good-outcomes patients were transplanted, and one of the transplanted patients died; see Figures 2A-B. In contrast, only 40% of HBC patients had good outcomes. Notably, like the 75% of older HBs, 78% of transplanted HBC patients had good outcomes. Finally, our observations confirmed previous reports based on smaller patient cohorts, which suggested that *TERT*-promoter mutations are of value as prognostic biomarkers (4). We established that these mutations are predominantly observed in cancers of older patients (Figure 2D) and may not be driver alterations.

Risk classification models—including CHIC-HS (14) and models based on molecular features by Cairo *et al*. (20) and Sumazin *et al*. (4)—suggested that nearly all patients in our cohort were high risk (Figure 2A), with the Cairo model suggesting that all tested patients are high-risk, and the Sumazin model agreeing for all but 3 patients. These predictions largely confirmed those based on a combination of demographic, surgical staging, and molecular features by CHIC-HS. In general, HBC risk stratifications were in good agreement with observed outcomes, however, 2 of the 3 HBC patients that were predicted to be low-risk by the Sumazin model were identified as high-risk by CHIC-HS and all 3 had good outcomes. Thus, the CHIC-HS risk stratification therapeutic group assignment may expose low-risk patients to unnecessarily aggressive and toxic therapy.

### HBC genomes are more genetically unstable

Other than recurring *CTNNB1* mutations or mutations in other WNT pathways genes, including *APC*—these features are observed in 90% of HBs—HBs have been reported to have relatively few coding mutations—at a total rate below 3-coding mutations per tumor (3, 12) or 0.2 somatic mutations per Mb of sequenced DNA (4). Tumor biopsies from our patient cohort—31 of which were profiled using the Texas Children’s Hospital Pediatric Solid Tumor (PST) panel—suggested a much higher mutation rate. The PST targets 124 genes and has a total capture size of 967-kilo bases. Our profiles revealed an average of over 3 coding mutations per biopsy, or a rate of 3.1 somatic mutations per Mb of sequenced DNA—a 16-fold increase in mutation rates relative to previous observations (4). This mutation rate approached the 3-5 somatic mutation per Mb rate observed for adult HCCs (21). While PST-targeted DNA is likely enriched for alterations, its relatively small target region identified as many alterations in our tumors as the expected total number of coding alterations in HBs (12). Moreover, while tumors of older HB patients are known to have higher somatic coding mutation rates (4), most patients under 8-years of age in our cohort had 3 or more coding mutations in PST-targeted regions; see Figure 3A.

**Figure 3.**
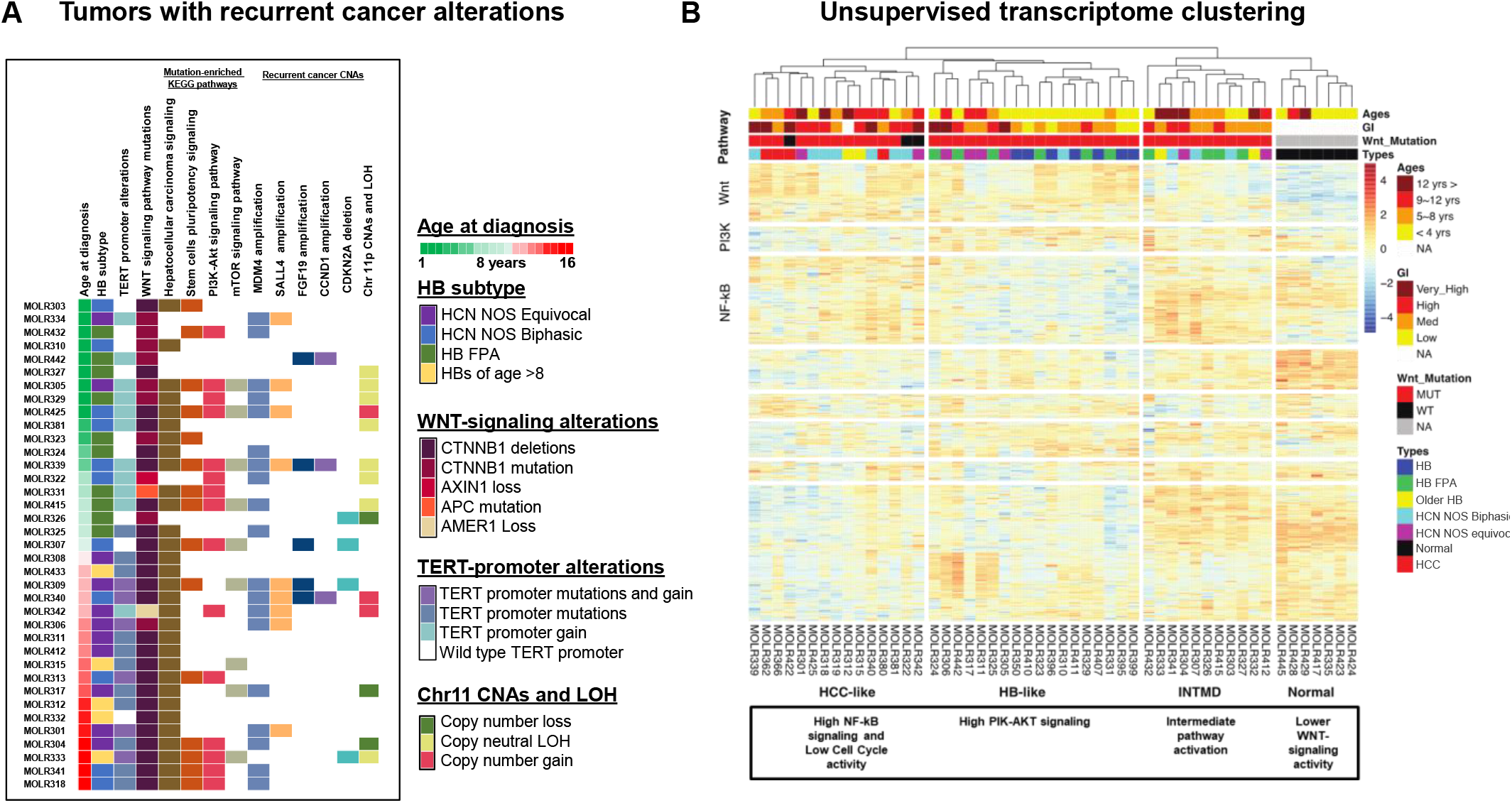
Unlike low-risk HBs, which harbor few genetic alterations outside of *CTNNB1*, our cohort revealed recurrent mutations, copy number alterations (CNAs), and dysregulation of multiple cancer genes and pathways. (**A**) Consistent with previous reports, all of our HBs had alterations in Wnt-signaling pathway genes, including *CTNNB1* and *APC*. In addition, however, we observed recurrent mutations and CNAs—defined as appearing in multiple patients—in genes and regions that are commonly altered in HCCs as well as genes that are associated with pluripotency signaling and common cancer pathways including PI3K-AKT and mTOR signaling. HBs were sorted by patient age at diagnosis; no significant associatiations between mutation rates or types with other tumor classifications were observed. Mutated genes in each pathway are given in Suplementary Table S2. (**B**) Unsupervised clustering of the expression profiles of non-cancer liver samples, HCCs, low-risk HBs, HCN NOSs, and HB FPAs revealed clusters that are enriched with dysregulated genes from at least 4 cancer pathways. These included, from left to right, clusters composed of (1) all profiled HCCs and some older, HB FPAs, and HCN NOSs; (2) all profiled low-risk HBs and some HB FPAs and HCN NOSs; (3) some older, HB FPAs, and HCN NOSs; and (4) non-tumor samples. Clusters 1-3 tumors had marked WNT-signaling pathway activation even in tumors with no identified activating mutations in Wnt-signaling pathway genes. Cluster 2 tumors had lower genetic instability (GI) and were seen in younger patients. Cluster 3 tumors had intermediate pathway activation compared to Cluster 1 tumors that showed high activation of NF-kB signaling and Cluster 2 tumors that showed high PIK-AKT signaling. Consequently, we labeled Cluster 1 HCC-like, Cluster 2 HB-like, Cluster-3 Intermediate (INTMD), and Cluster 4 Normal. Note that not all gene clusters were enriched for KEGG-pathway gene dysregulation.

Profiled tumors were enriched with mutations and alterations in key cancer genes and pathways. All patients had alterations in Wnt-signaling pathways, and, in addition, most patients had mutations in genes reported in HCC—even after excluding the commonly altered *CTNNB1—*and many patients had alterations in pathways that are associated with stem-cell pluripotency, PI3K-AKT, and mTOR signaling; see Figure 3A. The most common mutations were observed in the *TERT* promoter (51%), *FGFR4* (13%), *KMT2C* (10%), *KEAP1* (8%), *RPS6KA3* (8%), *CDK12* (8%), *NOTCH1* (8%), *BRCA2* (8%), *ARID1A* (5%), *ARID1B* (5%), *EP300* (5%), *MAPK1* (5%) and *PIK3CA* (5%); see Table S2 for a complete list. Amplifications targeting *MDM4*, *SALL4*, *FGF19*, and *CCND1*, as well as deletions targeting *CDKN2A* and *IRF2*, were also observed in multiple patients (Figure 3A). Confirming observations by Hirsch et al. (22), we identified recurrent copy number alterations and loss of heterozygosity (LOH) at chromosome 11p in nearly 40% of our samples. A complete map of copy number alterations (CNAs), including significant focal deletions and amplifications, is given in Figure 4 and Table S3. *TERT*-promoter, *KMT2C*, and *FGFR* mutations are hallmarks of aggressive cancers (23), while *KEAP1* mutations and *SALL4, CCND1*, and *CDKN2A* CNAs have been previously associated with poor outcomes for HB patients (4, 10, 24).

**Figure 4.**
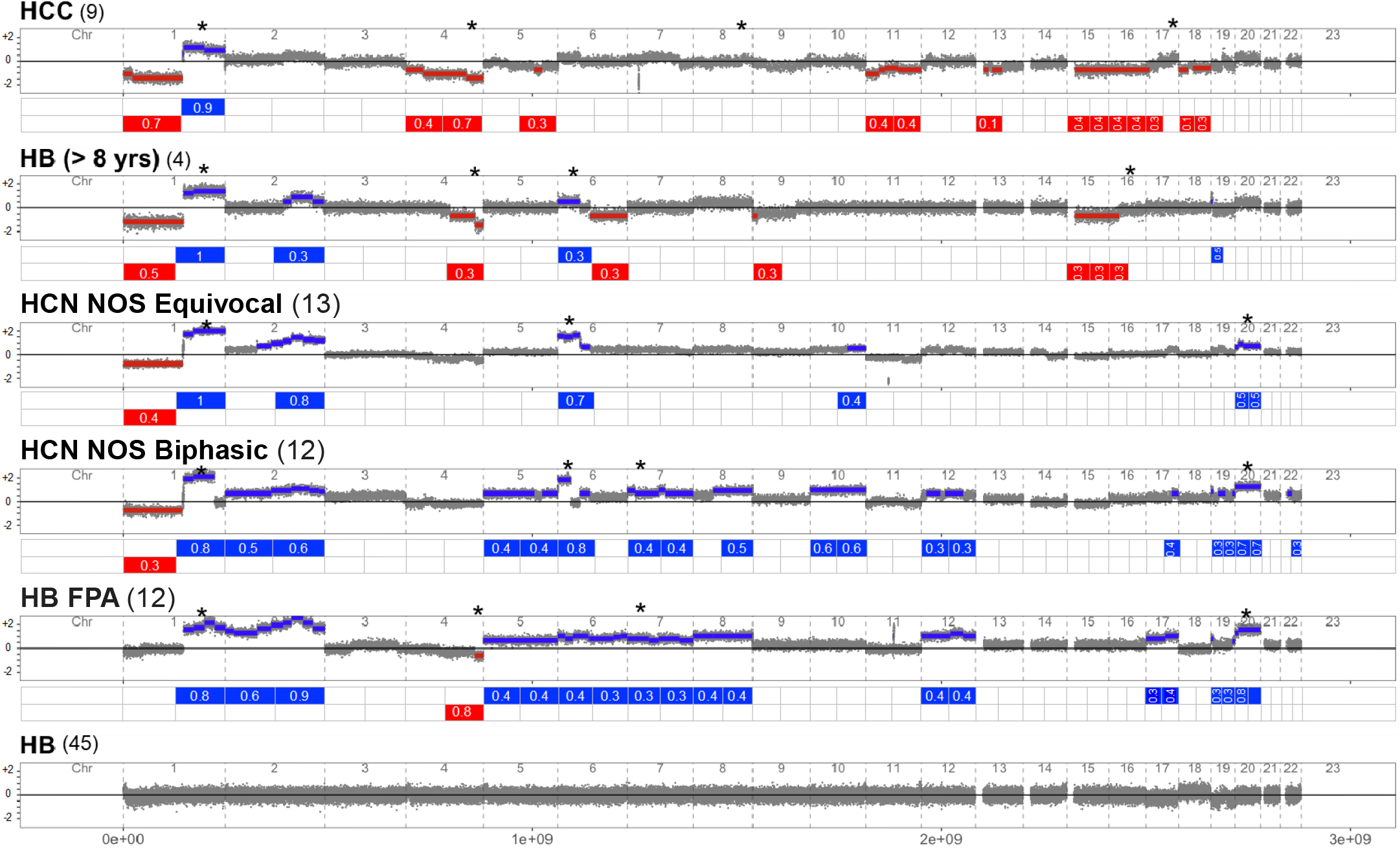
The copy number landscape of HCCs, older HBs, equivocal and biphasic HCN NOSs, HB FPAs, and low-risk HBs displaying common and differentiating patterns of whole-arm and focal amplifications and deletions. These included whole-arm amplifications of chr1q that were common to all HCCs and high-risk HB subtypes but not found in low-risk HBs; amplifications of chr2q that were typical of high-risk HBs; recurrent amplifications of chromosome 6p in HCN NOSs and some older HBs; and amplifications of chromosome 20p that were typical of HCN NOS and HB FPA patients. Deletions are known to be common in HCC and are present in some older HB patients, but, interestingly, chromosome 4q deletions are present in HCCs, HB FPAs diagnosed in younger children, and one of the HBs diagnosed in an older child. Lower risk HBs were retrieved from Sumazin et al. (2017) and included no significantly recurrent CNAs.

### HBCs showed common cancer gene and pathway dysregulation

We profiled gene expression in 37/40 (93%) of tumor samples using the NanoString nCounter assay, including 10 equivocal HCN NOS, 12 biphasic HCN NOS, 10 HB FPAs, and 5 HBs of older patients with usual HB histology. Unsupervised clustering of these profiles—together with control profiles of 7 low-risk HBs, 7 non-cancer pediatric liver samples, and 4 pediatric non-fibrolamellar HCCs with upregulated WNT-signaling pathway genes and no virus detection (6)—revealed marked differences between the profiles of cancer and non-cancer samples and similarities between the expression profiles of HBCs and low-risk HBs as well as of HCCs. Namely, as shown in Figure 3B, all non-cancer profiles clustered together, as did all low-risk HBs and all HCCs. However, while profiles of some HBCs and HBs of older patients clustered with HCCs, others clustered with HBs, and a third group formed an independent cluster. Interestingly, HB FPAs, which were selected in patients younger than 8 years, did not cluster with HCCs, while older HBs did not cluster with low-risk HBs. On the other hand, equivocal HCN NOSs, nearly all of which were collected from older patients, tended to cluster with low-risk HBs. Only one biphasic HCN NOS—MOLR310, who was younger than 2 at diagnosis—clustered with low-risk HBs.

Identified clusters were enriched with dysregulated genes from at least 4 cancer pathways. Tumor profiles showed upregulation of WNT-signaling pathway genes. The cluster containing low-risk HBs included cancers with upregulated PI3K-AKT signaling-pathway and cell-cycle genes, while the cluster containing HCCs included cancers with upregulated NF-kB signaling pathway genes. Interestingly the third cluster, composed of profiles of HBCs and older HBs that did not cluster with low-risk HBs or HCCs showed intermediate expression levels of these pathways; see Figure 5A. This category included 3 biphasic HCN NOSs, 2 equivocal HCN NOSs, 4 HB PFAs, and 2 older HBs. It also included patients from a variety of ages, including 3 and 4 patients diagnosed before the age of 4 and after the age of 12, respectively. Consequently, we named the four Figure 3B clusters HCC-like, HB-like, Intermediate (the third cluster), and Normal.

**Figure 5.**
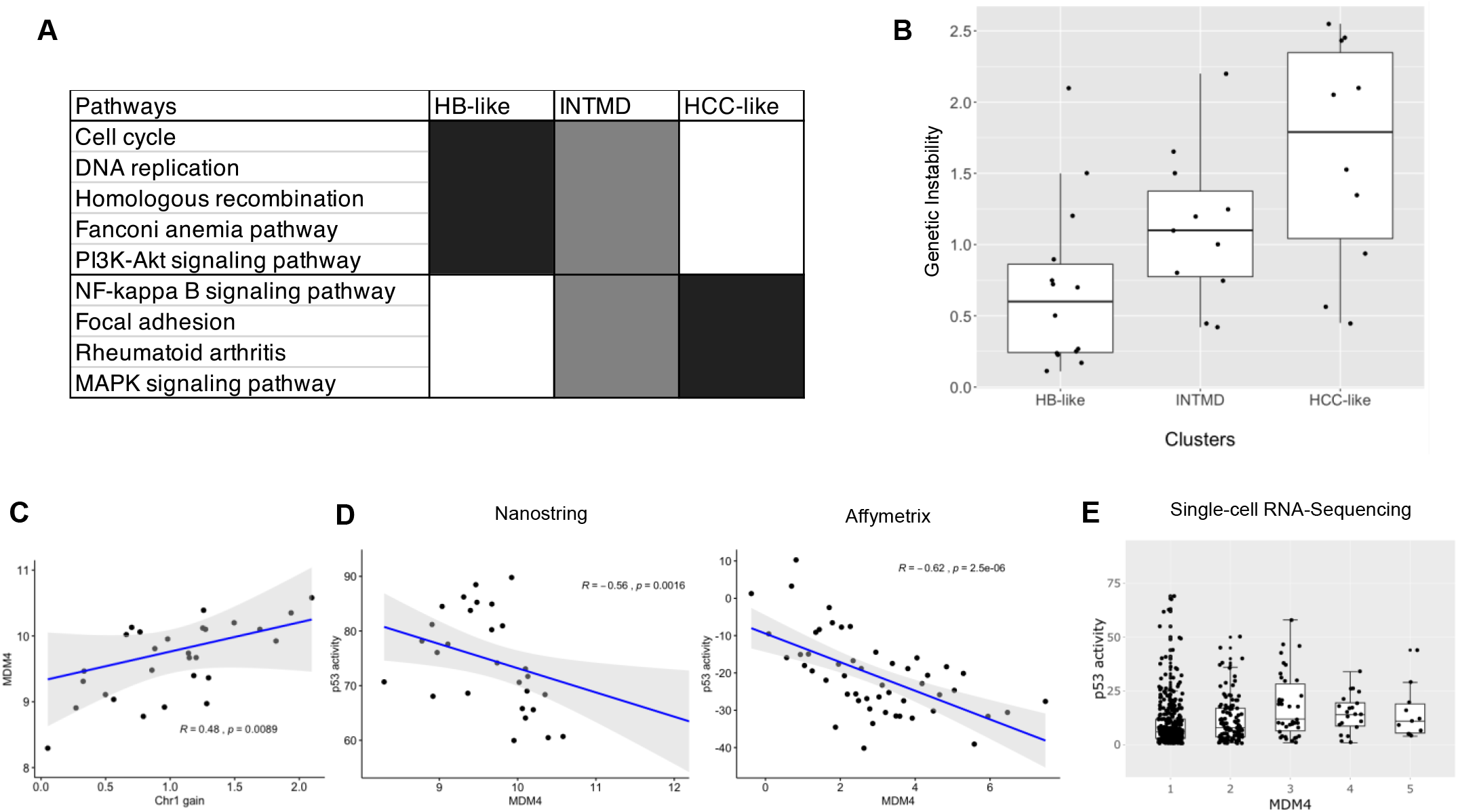
Genotypes and RNA-expression profiles of our tumors suggested that HCN NOSs and HB FPAs fall within a biological spectrum between low-risk HBs and HCCs and that their genetic instability may be associated with observed pathway dysregulation. (**A**) Our Intermediate (INTMD) expression cluster showed intermediate cancer-pathway activation compared to our HCC-like and HB-like clusters. (**B**) It also demonstrated an intermediate level of genomic instability—as calculated by the genome-wide CNA burden in each tumor—with HCC tumors having the highest genomic instability, on average. (**C**) High-risk tumors were enriched for chr1q amplification, which significantly coincided with higher MDM4 expression. (**D**) MDM4 expression profiles were inversely correlated with p53 activation in our dataset and in a second data set by Sumazin *et al*. (**E**) Single-cell profiling of a high-ris HB showed a significant correlation between MDM4 expression profiles and p53 activation in thousands of profiled cells.

### CNAs and genetic instability

HBs are known to generally have few CNAs and are genetically more stable than HCCs (4). To study the CNA landscape of our tumor set we profiled FFPE biopsies using the Affymetrix OncoScan assay. These SNP arrays detect CNAs at a 50kb resolution and were extensively validated in our laboratory for clinical use. To benchmark CNA detection in our set, we compared our profiles to profiles of 9 pediatric HCCs—including the 4 HCCs with upregulated WNT-signaling pathway genes—and 45 low-risk HBs (4). The full comparison is given in Figure 4. Our results confirmed that low-risk HBs have few CNAs and that HCCs have recurrent chromosome-1q amplifications and deletions in multiple chromosomes including 1p, 4, 11, 15, 16, and 18. HCCs also had recurrent focal deletions in chromosomes 8 and 17.

Profiles of HBs in older children demonstrated more frequent deletions than other HBs in the cohort, but all carried chromosome-1q amplification. The majority (90%) of HBCs had chromosome-1q amplification, but unlike older HBs and most profiled HCCs, we observed recurrent amplifications in these samples, including amplifications in chromosomal arms 2q, 6p, and 20p. We note that multiple amplified chromosomal regions detected matched those that were previously observed for WNT-pathway activated HCCs (6). Interestingly, chromosome 4q deletions were observed in HCCs, HB FPAs—in patients younger than 8 years—and some older HBs. We used CNAs to evaluate genetic instability at genome scales, scoring each tumor based on the number of nucleotides that were identified as amplified or deleted. Our analysis suggested that tumor samples identified as HB-like (Figure 3B) based on gene expression were the most genetically stable, while tumors that were identified as HCC-like were the most genetically unstable (Figure 5B).

Chromosome-1q amplifications were the most frequent genetic event in our high-risk cancer cohort, detected in nearly 90% of HBCs. Genes whose expression profiles were significantly correlated with their CNA profiles (p<0.01) included *THEM4, EFNA1, H3F3A, IL6R*, and *MDM4* (Figure 5C). Amplification of *MDM4* has been previously observed and associated with poor outcomes in HBs (25) and may inhibit p53 activity (26). Indeed, normalized expression of a panel of p53 target genes that have been studied in HCCs (21) and included in our NanoString platform—namely *CAPN2, CDKN1A, DUSP5, EPHA2, FAS, GADD45A, NFKBIA*, and *STAT3*—as a surrogate for p53 activity suggested that *MDM4* expression in both our and the Sumazin *et al*. datasets were significantly (p<0.01) anti-correlated with P53 activity (see Figure 5D). In addition, single-cell RNA sequencing of a frozen biopsy of the tumor MOLR442 showed that MDM4 RNA intra-tumor expression variability is also inversely correlated with p53 activity (Figure 5E).

## DISCUSSION AND CONCLUSIONS

Standardized diagnosis and histologic classification of pediatric hepatocellular neoplasms have been greatly aided by the development of the International Pediatric Liver Tumor Consensus Classification system (14). The provisional designation HCN NOS was proposed for a subset of difficult-to-classify neoplasms, acknowledging that further characterization would be required for cases in this provisional category. Therefore, our primary aim was to elucidate the histopathologic, molecular, genomic, and clinical features present in HCN NOSs. A database search to identify such cases also revealed the presence of HBs with atypical histological findings identified focally and most often post chemotherapy, and that did not meet the criteria for the HCN NOS diagnosis. Such findings include marked pleomorphism, anaplasia, and the presence of macro-trabecular patterns, as noted by the International Pediatric Liver Tumor Consensus Classification, and that are generally seen on post-therapy specimens.

Because of their similar histological and biological features and associated outcomes, we propose to designate three aggressive histological subtypes—including equivocal and biphasic HCN NOSs, and HB FPAs—as *HBs with carcinoma features* (HBCs). Our analyses of HBC biology suggested that these tumors share common molecular features with both HBs and HCCs and have recurrent genetic alterations that resemble WNT-pathway activated HCCs. Our retrospective analyses suggest that HBCs are high-risk tumors that may not respond to current HB treatment strategies (14) and that they can be histologically recognized and confirmed by molecular testing. Moreover, in our Texas Children’s Hospital dataset, HBCs that were treated by chemotherapy and surgery alone had significantly worse outcomes than transplanted HBCs, suggesting that these patients may benefit from aggressive and early interventions, including transplant referral when complete resection is not feasible. Based on our findings, we believe that this hypothesis merits further testing at a larger scale and in prospective trials that would account for surgical staging, clinical, histological, and molecular tumor features.

We argue that HBCs are more genetically unstable than lower-risk HBs and carry additional recurrent genetic alterations that target cancer genes (see Table 1). Our analysis suggested that WNT-driven hepatocellular tumors in children represent a biological and clinical spectrum and that HBCs are genetically distinct from classical lower-risk HBs, and share common features with Wnt-signaling driven pediatric HCCs. In addition, RNA-expression analyses suggested large-scale cancer-pathway dysregulation that may be associated with genetic pathogenic variants and copy number alterations. Our analyses, including the largest cohort of HCN NOSs analyzed to date, suggested that these cancers can afflict younger patients and that their biological hallmarks are recurrent genetic mutations and amplifications. Conclusive evidence for HBC’s associated clinical risk and recommendations for their treatment will only be achieved by identifying and characterizing this important group of tumors in upcoming international clinical trials, that should focus on incorporating molecular biomarkers to diagnose, risk stratify, and treat high-risk patients in general and HBCs in particular.

**Table 1.** Summary of biological features that are common to low-risk HBs, HB-HCCs, and HCCs.

|  | Low-risk hepatoblastoma (HB) | Hepatoblastoma with carcinoma features (HBC) | Hepatocellular carcinoma (HCC) |
| --- | --- | --- | --- |
| Mutation rates | 0.2 somatic mutations per Mb | 3 somatic mutations per Mb | 3-5 somatic mutations per Mb |
| Histology | Fetal, well-encapsulated, smaller cells | Both HB and HCC features, increased mitotic counts, higher nuclear-to-cytoplasmic ratios, macro-trabecular patterns, larger cells | High cellular density and nuclear-to-cytoplasmic ratios, larger cells, Cirrhosis |
| Recurrently altered cancer genes | CTNNB1 | CTNNB1, TERT promoter, FGFR4, KEAP1, KMT2C, MDM4, SALL4, FGF19, CCND1, CDKN2A, NFE2L2 | CTNNB1, FGF19, CCND1, VEGFA, TERT promoter, MYC, FAK, TP53, ARID1A, ARID2, AXIN, KEAP1, NFE2L2, RPS6KA3, ATM, RB1, RAS, MET, IGF2, PD1 |
| Pathways enriched for mutations | Wnt-signaling | Stem-cell pluripotency, Wnt, PIK, AKT/mTOR, and RAS/MAPK signaling; HCC pathway | Wnt, AKT/MTOR, and RAS/MAPK signaling; Telomere maintenance; Immune checkpoint |
| Recurrent copy number alterations | Rare 1q amplifications | 1q, 2q, 6p, 20 amplifications; 1p and 4q deletions | 1q amplifications; 1p, 4, 11, 15, 16, and 18 deletions; whole-arm amplifications genome-wide in WNT HCCs |

## ACKNOWLEDGMENTS

This work was funded by CPRIT award RP180674, the European Union’s Horizon 2020 research and innovation programme under grant agreement 826121, the Schindler Foundation, and NCI award R21CA223140. We thank the BCM GARP core for help in performing molecular assays. Raw data, including DNA and single-cell RNA sequencing, NanoString expression estimates, and OncoScan CNA predicitons will be made freely available from the European Nucleotide Archive, under study accession PRJEB46350.

---

**Figure S1.**
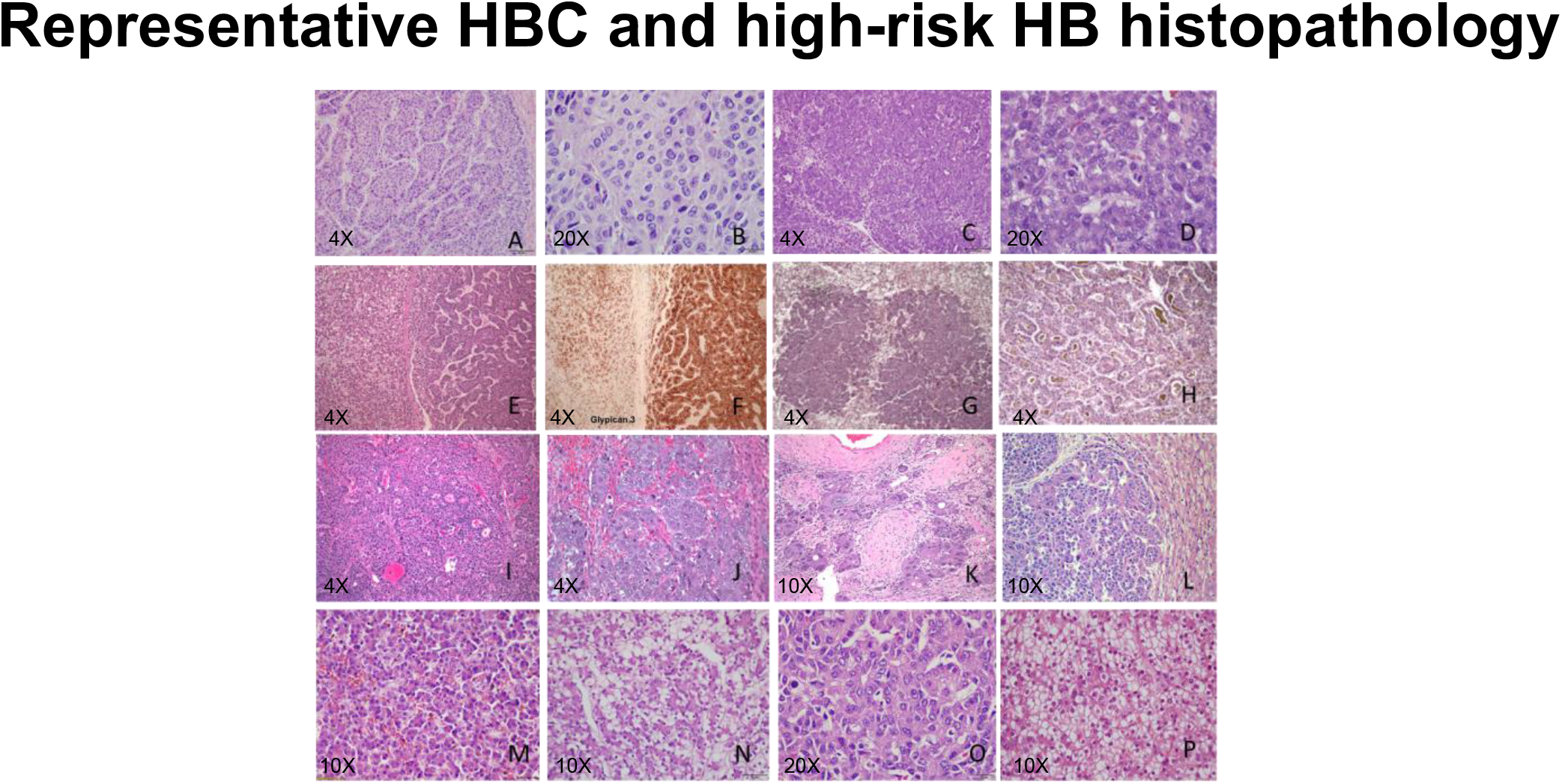
Histopathology of representative high-risk cases including (A, B) an equivocal HCN NOS with macrotrabecular growth patterns and (C, D) intermediate cytological features. (E, F) A biphasic HCN NOS with distinct areas demonstrating HB and HCC features and glypican 3 expression by immunohistochemistry. (G) A biphasic HCN NOS with distinct crowded fetal, and (H) HCC-like tubuloacinar bile producing areas. (I) An HCN NOS neoplasm with HB areas with (J-K) macrotrabecular and focal pleomorphic and anaplastic foci postchemotherapy, and (L) similar cytological features found in a metastasis. (M-O) Additional examples of HBs that were diagnosed in children older than 8 years of age, compared to a (P) fetal HB diagnosed in an 18-month-old child. Magnification levels (4X, 10X, or 20X) are posted in each figure.

**Figure S2.**
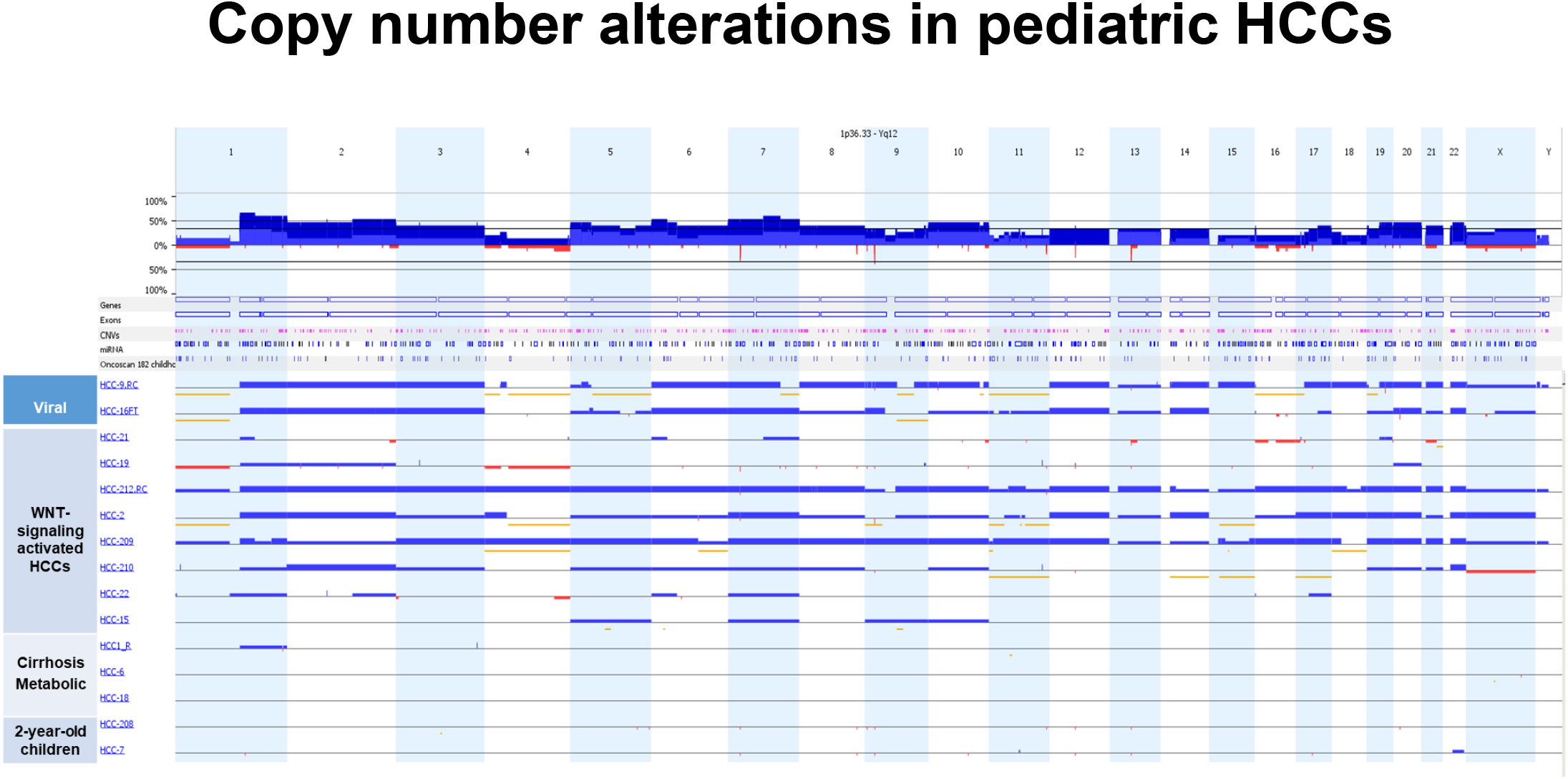
Visualization of whole-genome profiles by OncoScan of pediatric HCCs associated with viral infections (HCC-9 and 16), upregulated WNT-signaling pathway (HCC-21, 19, 212, 2, 209, 210, 22, and 15), and liver cirrhosis (HCC1, 6, and 18), compared to profiles of young patients under 2 years of age (HCC-208 and 7).

## REFERENCES

1. Hubbard AK, Spector LG, Fortuna G, Marcotte EL, Poynter JN. Trends in international incidence of pediatric cancers in children under 5 years of age: 1988–2012. JNCI cancer spectrum 2019;3:pkz007.

2. Czauderna P, Lopez-Terrada D, Hiyama E, Häberle B, Malogolowkin MH, Meyers RL. Hepatoblastoma state of the art: pathology, genetics, risk stratification, and chemotherapy. Current opinion in pediatrics 2014;26:19–28.

3. Ranganathan S, Lopez-Terrada D, Alaggio R. Hepatoblastoma and pediatric hepatocellular carcinoma: An update. Pediatric and Developmental Pathology 2020;23:79–95.

4. Sumazin P, Chen Y, Treviño LR, Sarabia SF, Hampton OA, Patel K, Mistretta TA, et al. Genomic analysis of hepatoblastoma identifies distinct molecular and prognostic subgroups. Hepatology 2017;65:104–121.

5. Ueda Y, Hiyama E, Kamimatsuse A, Kamei N, Ogura K, Sueda T. Wnt signaling and telomerase activation of hepatoblastoma: correlation with chemosensitivity and surgical resectability. Journal of pediatric surgery 2011;46:2221–2227.

6. Haines K, Sarabia SF, Alvarez KR, Tomlinson G, Vasudevan SA, Heczey AA, Roy A, et al. Characterization of pediatric hepatocellular carcinoma reveals genomic heterogeneity and diverse signaling pathway activation. Pediatric blood & cancer 2019;66:e27745.

7. López-Terrada D, Alaggio R, de Dávila MT, Czauderna P, Hiyama E, Katzenstein H, Leuschner I, et al. Towards an international pediatric liver tumor consensus classification: proceedings of the Los Angeles COG liver tumors symposium. Modern Pathology 2014;27:472–491.

8. Prokurat A, Kluge P, Kościesza A, Perek D, Kappeler A, Zimmermann A. Transitional liver cell tumors (TLCT) in older children and adolescents: a novel group of aggressive hepatic tumors expressing beta-catenin. Medical and pediatric oncology 2002;39:510–518.

9. Zimmermann A. The emerging family of hepatoblastoma tumours: from ontogenesis to oncogenesis. European Journal of Cancer 2005;41:1503–1514.

10. Eichenmuller M, Trippel F, Kreuder M, Beck A, Schwarzmayr T, Haberle B, Cairo S, et al. The genomic landscape of hepatoblastoma and their progenies with HCC-like features. J Hepatol 2014;61:1312–1320.

11. Carrillo-Reixach J, Torrens L, Simon-Coma M, Royo L, Domingo-Sàbat M, Abril-Fornaguera J, Akers N, et al. Epigenetic footprint enables molecular risk stratification of hepatoblastoma with clinical implications. Journal of hepatology 2020;73:328–341.

12. Cairo S, Armengol C, Maibach R, Häberle B, Becker K, Carrillo-Reixach J, Guettier C, et al. A combined clinical and biological risk classification improves prediction of outcome in hepatoblastoma patients. European Journal of Cancer 2020;141:30–39.

13. Haeberle B, Rangaswami A, Krailo M, Czauderna P, Hiyama E, Maibach R, Lopez-Terrada D, et al. The importance of age as prognostic factor for the outcome of patients with hepatoblastoma: Analysis from the Children’s Hepatic tumors International Collaboration (CHIC) database. Pediatric Blood & Cancer 2020:e28350.

14. Meyers RL, Maibach R, Hiyama E, Häberle B, Krailo M, Rangaswami A, Aronson DC, et al. Risk-stratified staging in paediatric hepatoblastoma: a unified analysis from the Children’s Hepatic tumors International Collaboration. The Lancet Oncology 2017;18:122–131.

15. Towbin AJ, Meyers RL, Woodley H, Miyazaki O, Weldon CB, Morland B, Hiyama E, et al. 2017 PRETEXT: radiologic staging system for primary hepatic malignancies of childhood revised for the Paediatric Hepatic International Tumour Trial (PHITT). Pediatric radiology 2018:1–19.

16. Roebuck DJ, Aronson D, Clapuyt P, Czauderna P, de Ville de Goyet J, Gauthier F, Mackinlay G, et al. 2005 PRETEXT: a revised staging system for primary malignant liver tumours of childhood developed by the SIOPEL group. Pediatr Radiol 2007;37:123-132; quiz 249-150.

17. Meyers RL, Tiao G, De Goyet JDV, Superina R, Aronson DC. Hepatoblastoma state of the art: pre-treatment extent of disease, surgical resection guidelines and the role of liver transplantation. Current opinion in pediatrics 2014;26:29–36.

18. Junco PT, Cano EM, Dore M, Gomez JJ, Galán AS, Vilanova-Sánchez A, Andres A, et al. Prognostic Factors for Liver Transplantation in Unresectable Hepatoblastoma. European Journal of Pediatric Surgery 2018.

19. Trobaugh-Lotrario AD, Meyers RL, O’Neill AF, Feusner JH. Unresectable hepatoblastoma: current perspectives. Hepatic medicine: evidence and research 2017;9:1.

20. Cairo S, Armengol C, De Reynies A, Wei Y, Thomas E, Renard CA, Goga A, et al. Hepatic stem-like phenotype and interplay of Wnt/beta-catenin and Myc signaling in aggressive childhood liver cancer. Cancer Cell 2008;14:471–484.

21. TCGA. Comprehensive and Integrative Genomic Characterization of Hepatocellular Carcinoma. Cell 2017;169:1327–1341. e1323.

22. Hirsch TZ, Pilet J, Morcrette G, Roehrig A, Monteiro BJ, Molina L, Bayard Q, et al. Integrated genomic analysis identifies driver genes and cisplatin-resistant progenitor phenotype in pediatric liver cancer. Cancer discovery 2021.

23. Zehir A, Benayed R, Shah RH, Syed A, Middha S, Kim HR, Srinivasan P, et al. Mutational landscape of metastatic cancer revealed from prospective clinical sequencing of 10,000 patients. Nature medicine 2017;23:703.

24. Iolascon A, Giordani L, Moretti A, Basso G, Borriello A, Della Ragione F. Analysis of CDKN2A, CDKN2B, CDKN2C, and cyclin Ds gene status in hepatoblastoma. Hepatology 1998;27:989–995.

25. Arai Y, Honda S, Haruta M, Kasai F, Fujiwara Y, Ohshima J, Sasaki F, et al. Genome-wide analysis of allelic imbalances reveals 4q deletions as a poor prognostic factor and MDM4 amplification at 1q32. 1 in hepatoblastoma. Genes, Chromosomes and Cancer 2010;49:596–609.

26. Woodfield SE, Shi Y, Patel RH, Chen Z, Shah AP, Whitlock RS, Ibarra AM, et al. MDM4 inhibition: a novel therapeutic strategy to reactivate p53 in hepatoblastoma. Scientific reports 2021;11:1–17.

